# Slow life-history strategies are associated with negligible actuarial senescence in western Palearctic salamanders

**DOI:** 10.1101/619494

**Authors:** Hugo Cayuela, Kurtuluş Olgun, Claudio Angelini, Nazan Üzüm, Olivier Peyronel, Claude Miaud, Aziz Avci, Jean-François Lemaitre, Benedikt R. Schmidt

**Author notes:** Corresponding author: Hugo Cayuela.

## Abstract

Actuarial senescence (hereafter “senescence”) has been viewed for a long time as an inevitable and uniform process. However, the work on senescence has mainly focused on endotherms (especially mammals) with deterministic growth and low regeneration capacity at adult stages, leading to a strong taxonomic bias in the study of aging. Recent studies have highlighted that senescence could indeed display highly variable trajectory shape that correlates with species life history traits. Slow life histories and indeterminate growth seem to be associated with weak and late senescence. Furthermore, a few studies have suggested that high regenerative abilities could make senescence negligible in several ectotherms (e.g., hydra and salamanders). However, demographic data for species that would allow testing of these hypotheses are scarce and fragmented. Here, we investigated senescence patterns in a group of salamanders (i.e. “true salamanders”) from the Western Palearctic using capture-recapture data and Bayesian modeling. Our results showed that salamanders have slow life histories and that they experience negligible senescence. This pattern was consistent at both intra- and interspecific levels, suggesting that the absence of senescence may be a phylogenetically conserved trait. The regenerative capacities of true salamanders, and urodeles in general, likely explains why these small ectotherms have lifespans similar to that of large endotherms (e.g., ungulates, large birds) and undergo negligible senescence contrary to most amniotes including humans. Our study seriously challenges the idea that senescence is a ubiquitous phenomenon in the living world.

## Introduction

The great diversity of life histories continues to fascinate population biologists (Stearns 1976, Dammhahn et al. 2018, Wright et al. 2019). There have been many attempts to summarize the variety of life history patterns that are observed (e.g., Pianka 1970, Sæther 1988, Bielby et al. 2007). Among the spate of life history traits, senescence is generally defined as a physiologically-caused, irreversible increase in mortality (i.e., actuarial senescence; hereafter senescence) and a decline in fertility with age (i.e., reproductive senescence) (Hamilton 1966, Rose 1991). Historically, senescence was expected to show a limited amount of variation across species. The senescence process was theoretically expected to be ubiquitous among age-structured populations (Hamilton 1966), which led to the view that senescence was an unavoidable process (Ackermann et al. 2003, Nussey et al. 2013). In addition, regardless of the species considered, the age at the onset of senescence was expected to be immutably set at the age of first reproduction (Williams 1957, Hamilton 1966). This long-held view was overturned when it was shown that the there is a bewildering diversity of senescence patterns across animal and plant species (Jones et al. 2014, Baudisch et al. 2013, Lemaître & Gaillard 2017, Colchero et al. 2019). There is an obvious need to further describe and explain the diversity of senescence patterns which was recently reinforced by the observation that senescence patterns can strongly affect population growth rates and how species respond to environmental change (Colchero et al. 2019).

In a recent analysis of senescence on a wide range of taxa Jones et al. (2014) demonstrated patterns of senescence are more diverse than previously thought. Although Jones et al. (2014), and even more recently Colchero et al. (2019), present results from a broad variety of species, their studies highlighted the lack of data for many taxa and thus there is still a great need to uncover and explain senescence patterns. In particular, it is necessary to evaluate the extent to which species can escape senescence or even show ‘negative senescence’ and to determinate the eco-evolutionary roots of such patterns (Vaupel et al. 2004, Jones & Vaupel 2017). As the Life History Theory of Aging postulates that senescence is related to somatic maintenance (Lemaître et al. 2015, Kirkwood 2017), one may expect that species who invest more into this item of expenditure such as “slow” species, (i.e., low recruitment and high adult survival), (see Ma & Gladyshev 2017) should show slower rates of senescence. This expectation is supported by the findings of Jones et al. (2008) who have shown that in both mammals and birds, the onset and rate of senescence are predicted by generation time and age at maturity. Since the life histories of species can be arranged along a fast-slow life history continuum (Harvey & Zammuto 1985, Promislow & Harvey 1990, Bielby et al. 2007), the speed of the life history can be used to predict senescence (Jones et al. 2008). Later, Jones et al. (2014) showed that the initial explanation of Jones et al. (2008) did not fully account for the diversity of senescence patterns, suggesting that other mechanisms may be at work as well. For example, their study revealed the absence of senescence in *Hydra* (and few other organisms), potentially allowed by regenerative capacity (Martinez 1998).

Most of our understanding of senescence in wild animal populations is based on mammals and birds because there are many long-term individual-based data sets (Jones et al. 2014). Yet, patterns of senescence among mammals is rather uniform. For example, Colchero et al. (2019) found that bathtub-shaped mortality trajectories were most commonly observed in ungulates and carnivores. The data set of Colchero et al. (2019) included only four amphibian species but four different models explained the data best (simple logistic, bathtub logistic, Gompertz, Weibull), suggesting great interspecific diversity within the amphibians. Interestingly, senescence was negligible in a salamander, an organism with an indeterminate (i.e., continuous) growth (Colchero & Schaible 2014, Jones & Vaupel 2017) that is well known for its regenerative capacity at the adult stage (Yokoyama 2008, Poss 2010, Seifert & Voss 2013).

Here, we investigated how slow life histories with an implicit high level of investment in somatic maintenance are associated with a negligible actuarial senescence in a clade of salamanders from the Western Palearctic (known as the “true salamanders”). We used both unpublished and published capture-recapture data for our analyses. First, we analyzed novel demographic data from a poorly known Mediterranean salamander (*Lyciasalamandra fazilae*) and verified that this species had a slow life history (i.e., high adult survival and low recruitment) consistent with the other species of true salamanders (Warburg 2007, Sinsch et al. 2017). Then, we examined senescence patterns in *L. fazilae* and two others species, *Salamandrina perspicillata* and *Salamandra Salamandra*, from western Europe. We expected negligible senescence in the three taxa and hypothesized that this pattern was consistent among populations.

## Material and methods

### *Demography of* Lyciasalamandra fazilae

#### Study species and capture-recapture survey

*L. fazilae* is terrestrial salamander occurring along the southern Anatolian coast in Turkey. This species is a member of the phylogenetic clade called “true” salamanders that encompasses the genera *Salamandra, Lyciasalamandra, Mertensiella, Salamandrina*, and *Chioglossa* (Weisrock et al. 2006). *L. fazilae* is viviparous salamander that gives birth to one or two fully metamorphosed young after one year of gestation. Sexual maturity is attained at an age of three years in both sexes (Olgun et al. 2001). The individual growth curve presents an asymptotic trend even if adult salamanders seem to continue to grow over their entire lifespan (Olgun et al. 2001).

The study was conducted on a population of *L. fazilae* between 1999 and 2009 in western Turkey near Dalyan (N 36° 50’, E 28° 41’). A detailed description of the study area can be found in Olgun et al. (2001). Several capture–recapture sessions (in February, March, and April) were carried out each year. The salamanders were captured by hand, and were then released back the place where they were initially caught after being marked using pit-tags, identified, and measured. The sex of the individuals was assessed using secondary sexual characters (Olgun et al. 2001). Juveniles were not included in the analysis because only 12 juveniles were encountered during the study but never recaptured. Furthermore, due to small size of the dataset (133 individuals marked), we grouped the two sexes in analyses to avoid model overparameterization. We assumed that sex should have a little influence on adult survival as males and females present the same age structure in the population (Olgun et al. 2001).

Body size of salamanders was determined by measuring snout-vent-length (SVL): individuals were measured from the tip of the snout to the posterior margin of the vent. We used the size data in multievent models to estimate size-dependent survival. We also benefited from age data assessed using skeletochronological analyses for individuals marked over the period 1999-2003 (Olgun et al. 2001). Those age data were included in Colchero’s models (Colchero et al. 2012a, 2012b) to examine age-dependent survival and mortality rate.

#### Multievent model with for size-dependent adult survival

We quantified size-dependent annual survival using multievent capture-recapture models (Pradel 2005). Note that we did not performed Goodness-of-fit test before building the models since no test is currently available for multievent models; note that it was also possible to consider potential transience, trap-dependence, and recapture heterogeneity in Colchero’s models presented below. We considered a model based on three latent states that include information about individuals’ size. The states *s* and *l* correspond to small (SVL, from 45 to 60 mm) and large (SVL, from 61 to 81 mm) adults respectively; the classes were fixed to obtain a relatively similar number of observations in the two classes. The state *d* corresponds to the *dead* state. The models include three observations coded as following in the capture histories: individuals that are not captured are coded ‘0’; small and large individuals captured are coded ‘1’ and ‘2’ respectively.

At their first capture individuals may occupy two distinct states of departure, s and *l*. At each time step, the information about individual state is progressively updated through two successive modeling steps: (1) survival, and (2) size transition. Each step is conditional on all previous steps. At the first modeling step, survival information is updated. A small individual may survive with a probability φ*_s_* or die with a probability 1–φ*_s_*. A large individual can survive with a probability φ*l* or not with a probability 1–φ_*l*_. These results in the following matrix (the state of the individual at *t*-1 is in column and state at *t* is in row):

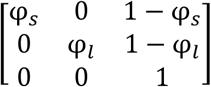

In the second modeling step, information about individual age is updated. A small individual can become a large individual with a probability *δ_s_* or remain in the same class with a probability 1– *δ_s_*, leading to the following matrix:

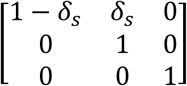

The last component of the model links observation to states. Small and large individual can be captured with a probability *p_s_* or *p_l_*, which results in the following matrix:

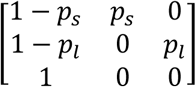

This parameterization was implemented in the program E-SURGE (Choquet et al. 2009). We ranked models using AICc and Akaike weights (w). If the Akaike weight of the best supported model was less than 0.9, we used model-averaging to obtain parameter estimates. We examined our hypotheses about survival and recapture probability from the following general model [φ(size), *δ*(.), *p*(t + size)]. The effects considered in the models were size and year (*t*). We hypothesized that: survival φ probability differed between the two size classes (i.e. small and large). We also expected that recapture probability varies according to size and year. Age transition was set constant (.) in the model. We tested all the possible combinations of effects, resulting in the consideration of eight competing models (**Table 1**).

**Table 1.**
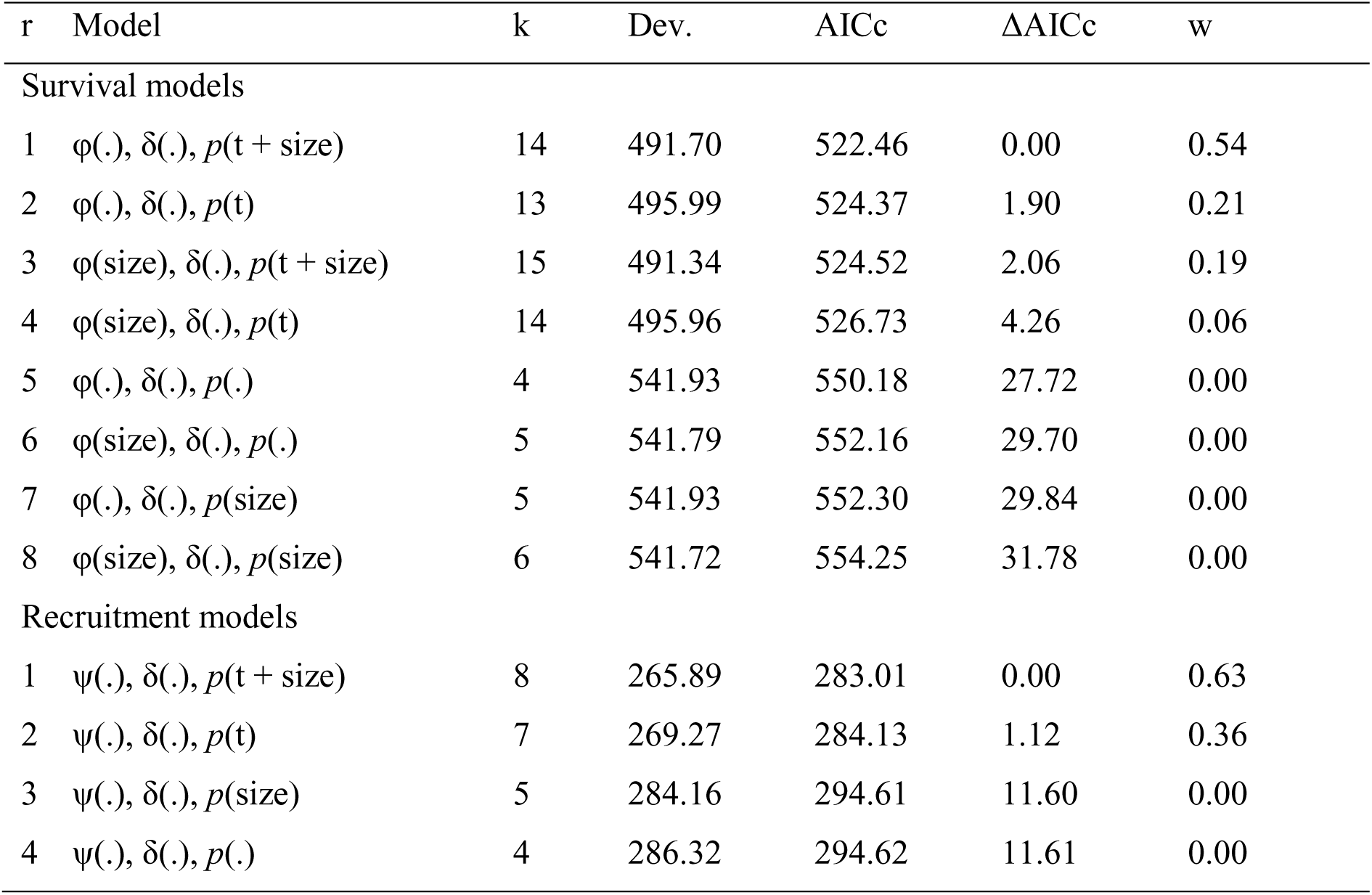
Multievent survival and recruitment models: model selection procedure. r = model rank, k = number of parameters, Dev. = residual deviance, AICc = Akaike Information Criterion adjusted for small sample size, δAICc = difference of AICc points with the best-supported model, w = AICc weight.

#### Multievent model for recruitment

We estimated recruitment rate of small-size adult using a modified Pradel model (1996) in which recruitment is modeled by reversing capture histories and analyzing them backwards. Recruitment probability was estimated as the probability that a small-sized individual present at *t* was not present at *t*-1, i.e. the proportion of “new” small individuals in the population at *t*. The model had the structure similar to that of survival model. The survival matrix was replaced by the recruitment matrix. At each time step, small individuals may be recruited with a probability *ψ*_s_ or not with a probability 1 – *ψ*_s_, leading to the following matrix:

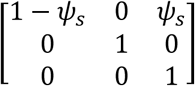

The size transition matrix was also modified to allow reversed size transition:

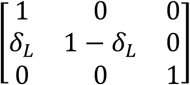

This parameterization was implemented in E-SURGE program. We considered the most general model [*ψ*(.), *δ*(.), *p*(t + size)]. We examined all the possible combination of effects leading to the consideration of four candidate models.

### Age-dependent survival and senescence patterns in true salamanders

#### Capture-recapture data

To estimate age-dependent survival and senescence, we re-analysed two data sets of true salamanders that were analysed previously. The capture-recapture data of *S. perspicillata* was collected over a 9-year period (1998-2006) in a population of central Italy, Monti Lepini, Latium (Angelini et al. 2010). The capture-recapture data of *S. salamandra* were collected in two populations from southern France (Ardèche region, *pop1*) and northwestern Germany (Nordrhein-Westfalen, *pop2*). *Pop1* and *pop2* were surveyed over 8-years (2008-2015) and 21-years (1965-1985) period, respectively (Cayuela et al. 2017, Schmidt et al. 2005). A summary of the age-dependent capture-recapture data in the four populations of salamanders is provided in Supplementary material, Table S1. Note that the sex was not considered in the further analyses as only females were caught in *S. perspicillata* (males do not occur in breeding sites) and because sex cannot be easily ascertained by non-expert observers in *S. Salamandra*.

#### Age-dependent survival and mortality rate

We investigated actuarial senescence patterns in the three salamander species using Bayesian survival trajectory analyses implemented in the R package BaSTA (Colchero et al. 2012a, 2012b). BaSTA allowed us to account for imperfect detection, left-truncated (i.e., unknown birth date) and right-censored (i.e., unknown death date) capture-recapture data in our analysis. It allows estimation of two parameters: survival until a given age and mortality rate (i.e., hazard rate) at a given age. Given the results of previous analyses (Schmidt et al. 2005, Angelini et al. 2010, Cayuela et al. 2017), we allowed recapture probabilities to vary among years. As the study period and number of survey years differ among populations (Supplementary material, Table S1), the four populations and species were analyzed separately. We used DIC to select models that fit the data best and we compared the outputs of the best-supported model of the four populations by inspecting mean estimates and 95% CI. This allowed us to investigate population/species-specific variation in the shape of the age-specific mortality patterns. We considered the four mortality functions implemented in BaSTA: exponential, Gompertz, Weibull and logistic. For the three last functions, we considered three potential shapes: *simple* that only uses the basic functions described above; *Makeham* (Pletcher 1999); and *bathtub* (Siler 1979). As individuals cannot be individually surveyed before their sexual maturity (three years old in the three species), we conditioned the analyses at a minimum age of three. Four MCMC chains were run with 50000 iterations and a burn-in of 5000. Chains were thinned by a factor of 50. Model convergence was evaluated using the diagnostic analyses implemented in BaSTA, which calculate the potential scale reduction for each parameter to assess convergence. For all populations, we used DIC to compare the predictive power of each mortality function and its refinements (Spiegelhalter et al. 2002, Colchero et al. 2012b).

## Results

### *Demography of* Lyciasalamandra fazilae

Over the 8-year study period, we made 179 captures of salamanders. We identified 121 adults (51 males and 70 females) and 12 juveniles. The mean age was 5.5 years and the maximum was 10 years.

The best-supported survival model was [φ(.), *δ*(.), *p*(t + size)] (**Table 1**); its AICc weight was 0.54 and we therefore model-averaged the estimates. The recapture probability of small individuals was higher than that of large individuals and varied over time. In 2001, the recapture probability of small individuals was 0.05±0.05 while it was 0.01±0.01 in large individuals; in 2007, recapture probability of small and large individuals was 0.78±0.18 and 0.41±0.17 respectively. The probability that a small individual change of size class and became a large individual on was 0.28±0.09. Survival did not differ between small (0.72±0.06) and large (0.75±0.04) individuals (**Fig.2A**).

**Fig 1.**
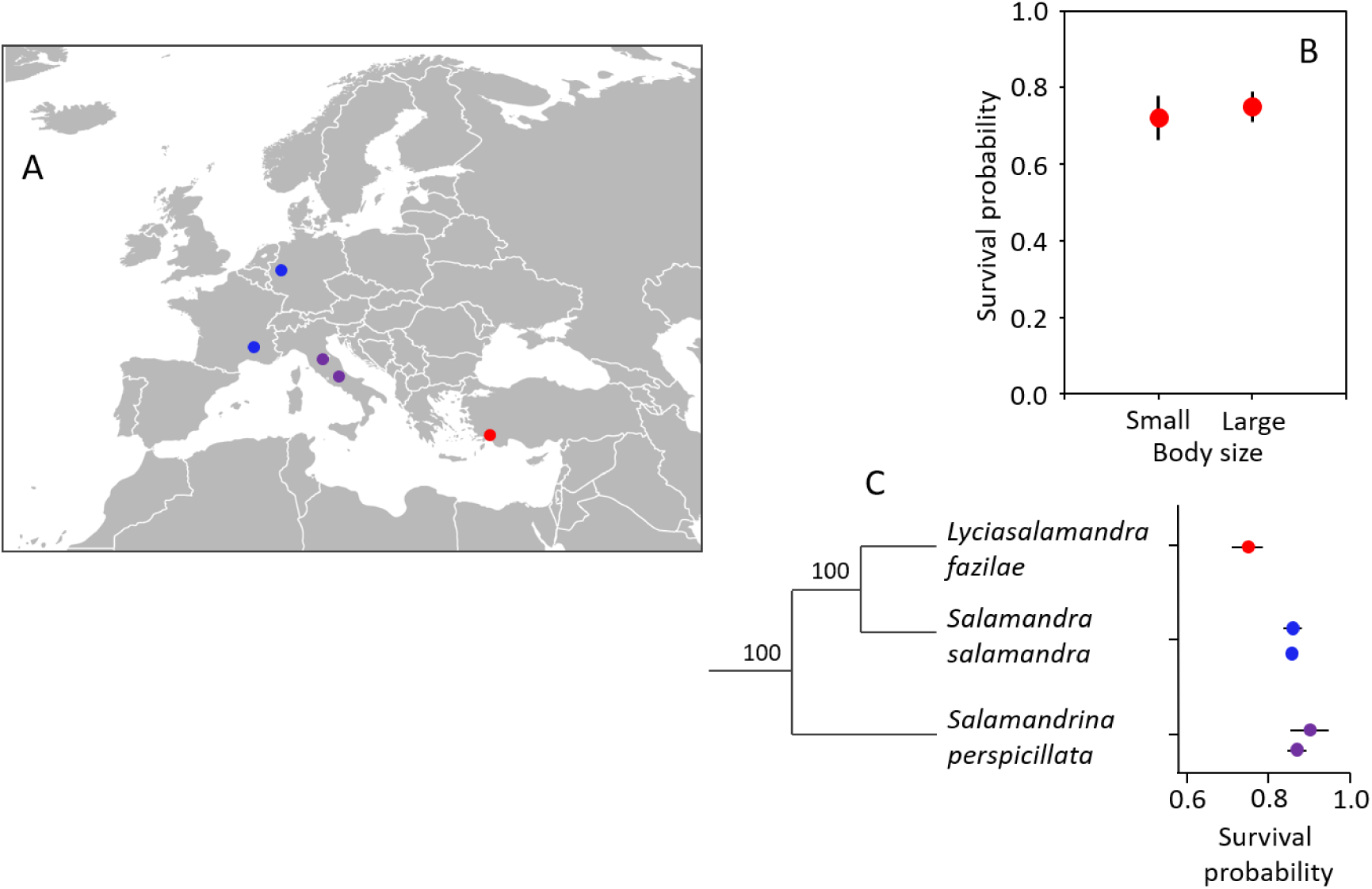
Adult survival (and standard errors) in three species of salamanders in western Palearctic. (A) Map showing the populations of *Lyciasalamandra fazilae* (in red), *Salamandra salamandra* (in blue), and *Salamandra perspicillata* (in violet) considered in this study. Note that data of only one population of *Salamandra perspicillata* was available for the present study (the southernmost population on the map). (B) Survival of small and large adults of *Lyciasalamandra fazilae* and their standard errors (model-averaged) estimated using our survival and recruitment models. (C) Survival estimates of *Lyciasalamandra fazilae* (from this study) and *S. salamandra* and *S. perspicillata*, which are extracted from two previous capture-recapture studies; i.e., Cayuela et al. (2017) and Angelini et al. (2010) respectively. The phylogenetic tree is the one presented in Weisrock et al. 2006.

**Fig 2.**
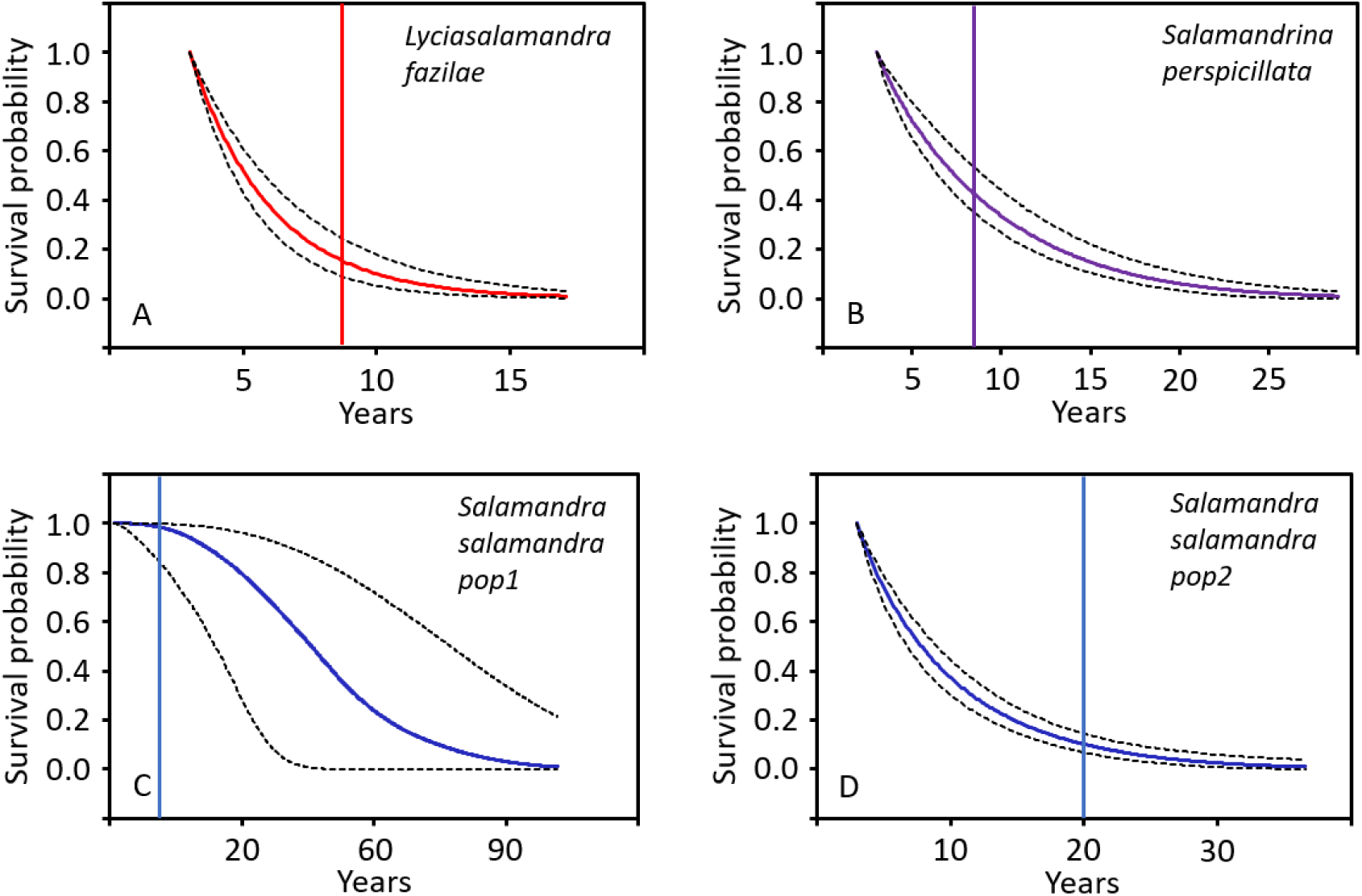
Survival until age *x* (full lines) and their 95% CI (broken lines) in three salamanders of western Palearctic. The vertical line indicates the length of the capture-recapture survey.

The best-supported recruitment model was [*ψ*(.), *δ*(.), *p*(t + size)] (**Table 2**); its AICc weight was 0.54 and we therefore model-averaged the estimates. The recapture probabilities were relatively similar to that provided by survival model; we did not report them for this reason. The rate of small individual recruitment was 0.15±0.15.

**Table 2.** Deviance information criterion (DIC) for each of the mortality function in three salamanders of western Paleartic. We considered the four mortality functions implemented in BaSTA program: exponential (EXP), Gompertz (GOM), Weibull (WEI) and logistic (LOG). For the three last functions, we considered three potential shapes: simple that only uses the basic functions described above (“Simple”); Makeham (“Makeham”); and bathtub (“Bath”).

|  | *Simple* | *Bath* | *Makeham* |
| --- | --- | --- | --- |
| *Lyciasalamandra fazilae* |  |  |  |
| Logistic | Not converged | Not converged | 1909.03 |
| Exponential | **1593.92** | - | - |
| Gompertz | 1626.67 | Not converged | 1946.88 |
| Weibull | 1709.25 | Not converged | 1701.72 |
| *Salamandrina perspicillata* |  |  |  |
| Logistic | 15394.47 | 14928.48 | 15382.08 |
| Exponential | Not converged | - | - |
| Gompertz | Not converged | **14881.98** | Not converged |
| Weibull | 15530.53 | 14898.80 | 15238.54 |
| *Salamandra Salamandra (pop1)* |  |  |  |
| Logistic | 6704.18 | Not converged | 6723.45 |
| Exponential | Not converged | - | - |
| Gompertz | 6479.37 | Not converged | Not converged |
| Weibull | **6319.93** | Not converged | 6711.14 |
| *Salamandra Salamandra (pop2)* |  |  |  |
| Logistic | 6099.82 | 6108.51 | 6101.58 |
| Exponential | 6137.15 | - | - |
| Gompertz | 6161.98 | **6045.60** | 6151.98 |
| Weibull | 6067.80 | 6111.62 | 6080.74 |

### Age-dependent survival and senescence in true salamanders

In *L. fazilae*, the age-specific capture-recapture data were best described by an exponential function (**Table 2**). The probability of surviving was 0.75 until age four, 0.50 until age five, and 0.25 until age seven (**Fig.2A**). Furthermore, the model indicates the absence of age-dependent mortality rate: mortality rate remained stable (around 0.34) regardless of age (**Fig.3A**).

**Fig 3.**
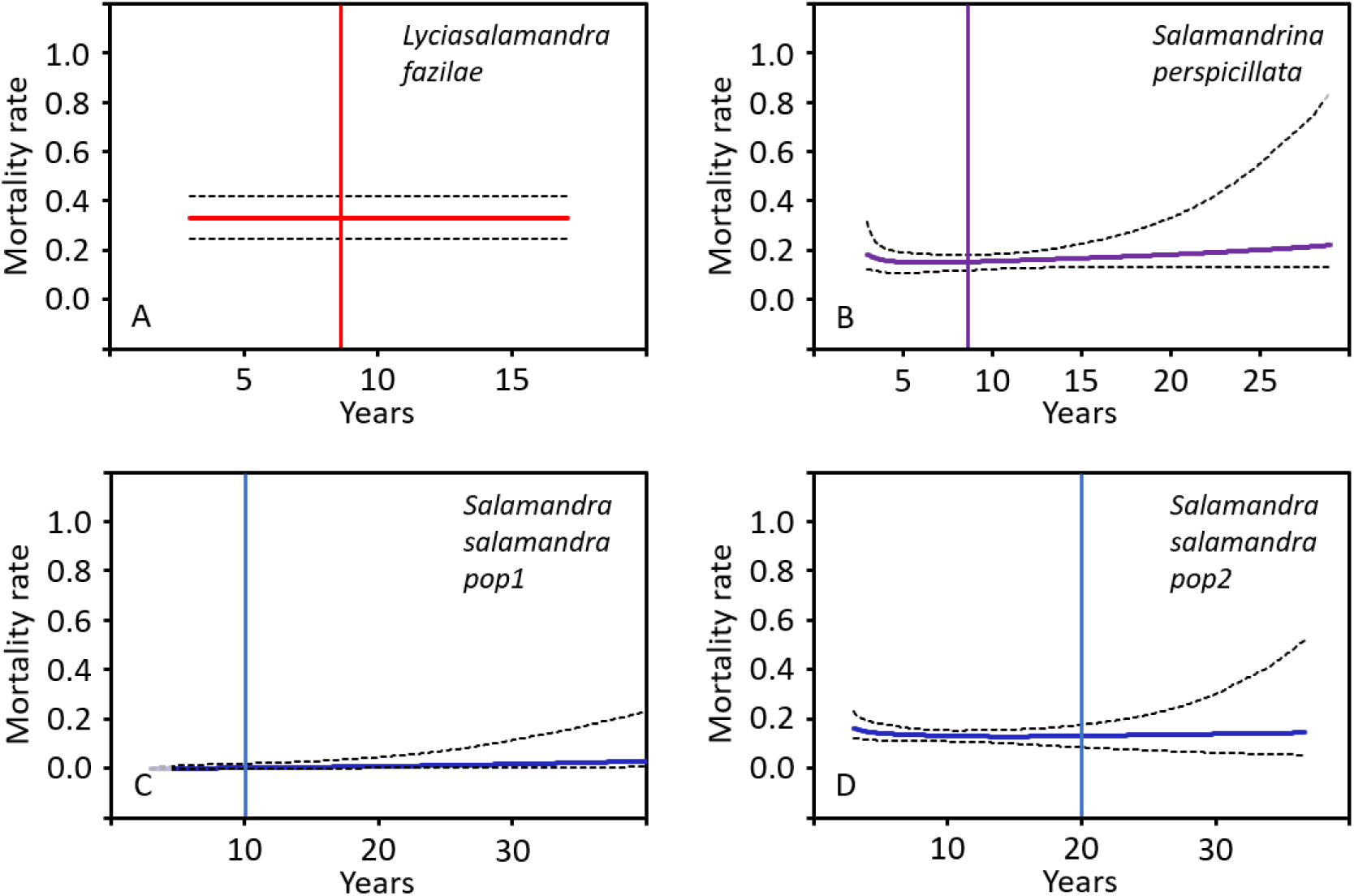
Mortality rate at age *x* (full lines) and their 95% CI (broken lines) in three salamanders of western Palearctic. The vertical line indicates the length of the capture-recapture survey.

In *S. perspicillata*, the best-supported model included a Gompertz function (**Table 2**). The probability of surviving was 0.75 until age five, 0.50 until age eight, and 0.25 until age 12 (**Fig.2B**). Moreover, our results indicate that age had no influence on mortality rate (**Fig.3B**). Mortality rate remained stable (around 0.20) regardless of age.

In *S. salamandra*, the best-supported model included a Weibull function in *pop1* and a Gompertz function in *pop2* (**Table 2**). The probability of surviving was 0.75 until age five, 0.50 until age eight, and 0.25 until age 13 in *pop2* (**Fig.2C**); in *pop1*, the survival estimates were very imprecise (**Fig.2D**). Moreover, our results indicate that mortality rate was not affected by age (**Fig.3C** and **3D**). Mortality rate remained stable (around 0.05 and 0.15 in *pop1*and *pop2* respectively) regardless of age.

## Discussion

The three species of true salamanders analysed in this study are characterized by a slow life history strategy (i.e., low recruitment and high adult survival). Survival of all three species of salamanders decreased slowly with age and mortality rate remained constant regardless of salamander age, suggesting negligible actuarial senescence in this clade.

### *Demography of* Lyciasalamandra fazilae

Using capture-recapture data we have provided the first detailed demographic characterization of L. fazilae and demonstrated that it is a species with a relatively “slow” life history. The survival estimate of our multievent model (mean adult survival = 0.74±0.05) was very close to the survival estimate (0.79) provided by the skeletochronological analysis in the same population (Olgun et al. 2001). We are therefore confident that the survival estimate of *L. fazilae* is not biased by a methodological artefact; for example, permanent emigration leads to underestimated survival rates (i.e., apparent survival) in capture-recapture studies (Schmidt et al. 2007). Our study revealed that the probability of surviving was 0.50 until age five, and less than 0.05 until age 12. These results are also congruent with skeletochronological data that reported a maximum longevity of 10 years in the population. Furthermore, we did not find evidence of an effect of body size on adult survival. It is possible that a small body size negatively affects survival at juvenile stage (as commonly reported in amphibians; e.g., Schmidt et al. 2012, Cayuela et al. 2016) and that the effect of size on survival vanishes at either some point before or rapidly after sexual maturation. Furthermore, our study also revealed that recruitment was relatively low in *L. fazilae* (0.15). This is likely due to the low fecundity of females that produce one or two young after a gestation of one year (Olgun et al. 2001). All together, these results indicate that *L. fazilae* has a relatively slow life history.

### Slow life histories in true salamanders

Our study and previous ones (Angelini et al. 2010, Cayuela et al. 2017) indicate that true salamanders have slow life history strategies. First, adult survival is relatively high in this clade (Fig.1C). In *S. Salamandra*, survival probability was 0.85 in both populations while it ranges from 0.86 to 0.90 *S. perspicillata*, indicating little variation at the intraspecific level (Fig.1C). At the interspecific scale, survival is relatively high in these three genera, which suggests that a long lifespan is a highly conserved trait in salamanders of western Palearctic. Yet, *L. fazilae* has a lower survival (0.74 and 0.79 from capture-recapture and skelettochronological analyses respectively) than *S. salamandra* and *S. perspicillata* (Fig.1C). This pattern was also perceptible in the survival estimates provided by our study: a survival probability of 0.50 was reached at 5 years in *L. fazilae*, 7.5 years in *S. perspicillata*, and 8 years in *S. Salamandra* (in *pop2*, the estimate of *pop1* was very imprecise). These adult survival rates for true salamanders are higher than for most anurans (Muths et al. 2017) and many newts and Plethodontids salamanders (Jehle et al. 1995, Bendik 2017, Cayuela et al. 2019) supporting the idea that these species exhibit slow life histories.

Through a trade-off, the relatively long lifespan of salamanders is also associated with a low fecundity (compared to other amphibians; Wells 2010, Oliveira et al. 2017) that is modulated by their reproductive modes. Oviparous species have the highest fecundity (e.g., *Salamandra infraimmaculata*, 218 eggs per female per year, Sharon et al. 1997; *S. perspicillata*, 40-65 eggs, Angelini 2006, Rovelli et al. 2015). Salamanders with lecithotrophic viviparity (i.e., giving birth to larvae) have an intermediate fecundity (e.g., *S. salamandra*, 23 larvae per female per year, Kopp & Baur 2000; *Salamandra algira*, 13 larvae, Reinhard et al. 2015) whereas species with matrotrophic viviparity (i.e., giving birth to fully developed young) have the lowest fecundity (e.g., *L. fazilae* and *Salamandra atra*, 1 young per female per year; Greven & Thiesmeier 1994, Luiselli et al. 2001). Fecundity adjustments of true salamander result from potentially rapid (less than 1000 generations, Velo-Antón et al. 2012) changes in reproductive modes that may occur at the intraspecific level (Velo-Antón et al. 2012). These characteristics make “true” salamanders an exceptional biological system to extend our understanding of the evolution reproductive modes and the modularity of life history strategies in amniotes.

### Negligible actuarial senescence in true salamanders

We observed negligible actuarial senescence in the three species of true salamanders considered in our study. The mortality rate was relatively stable and weakly affected by age in the three taxa and the two populations of *S. salamandra*. This mortality pattern markedly differs from age-dependent mortality rates found in various species of anurans and newts (using capture-recapture data with relatively similar sample size, and a similar modeling approach), which may experience a sharp, early senescence (Miller et al. 2014, Colchero et al. 2019; but also see Jones et al. 2014). The detection of actuarial senescence in other species using similar modeling tools (Miller et al. 2014, Colchero et al. 2019) indicates that the negligible senescence in true salamanders is not a methodological artefact. It has previously been suggested that detecting senescence would be difficult in wild animals because high levels of mortality would remove individuals from the population before they start to senesce (Kirkwood & Austad 2000). However, by revealing a great diversity of senescence patterns, studies using capture-recapture data collected across a broad range of taxa showed that this assumption was untrue (Jones et al. 2008, 2014; Colchero et al. 2019; see also the review of Jones & Vaupel 2017). Furthermore, Colchero’s model has proven to be particularly efficient for detecting senescence when it is actually present (Colchero et al. 2012a, 2012b). Simulations showed that the model is able to detect senescence when the study period is equal or longer than the mean lifespan in the population (Colchero et al. 2012a), which was the case in the four datasets considered in our study.

A long lifespan, a high level of iteroparity, and a low fecundity appear to be closely associated with negligible actuarial senescence in true salamanders. This pattern is congruent with the results of Jones et al. (2008) showing that senescence rate is negatively correlated with generation time and age at primiparity, and is positively associated with maximum fecundity in endotherm vertebrates; the opposite relationships were detected with the age at senescence onset. Contrary to other amniotes, salamanders have high regenerative capacities and are able to retain near perfect regeneration of most organs and appendages (e.g., spinal cord, heart, brain, skin, digit, and lens) well into adulthood (Seifert & Voss 2013). Although almost no studies have tested these abilities in old animals (Seifert & Voss 2013), their great potential for tissue repair and regeneration likely allow true salamanders to experience negligible actuarial senescence. In parallel, an indeterminate growth could also contribute to this marginal senescence, as proposed by Jones & Vaupel (2017).

Our study also showed that negligible senescence is a consistent pattern at both intraspecific and interspecific levels in true salamanders (at least in the species considered in this study). This indicates marginal senescence can be a phylogenetically conserved trait within a clade containing species that have diverged a long time ago (several million years; Weisrock et al. 2006). These results suggest a strong genetic determinism in aging mechanisms and the existence of orthologous genes involved in the repression of actuarial senescence in urodeles. However, the genomic architecture of life history components as senescence rate and onsets remains poorly understood, except in few model species (e.g., drosophila: Remolina et al. 2012, Ivanov et al. 2015; human: Deelen et al. 2013). The recent development of powerful genomic tools should allow to identify candidate genes and gene networks involved in senescence regulation.

## Conclusion

Negligible actuarial senescence is highlighted in a growing number of taxa, mainly ectotherms (e.g., corals, hydras, and amphibians; Jones et al. 2014). These cases have been considered for a long time as exceptions or the product of methodological artefacts, in light of senescence having been presented as a nearly ubiquitous phenomenon in the living world. We argue that this representation was partly due to a taxonomic bias where the study of senescence has for many years been focused on endotherm vertebrates (mainly mammals) with reduced regenerative capacities at adult stages (Seifert & Voss 2013). The regenerative capacities of true salamanders, and urodeles in general, likely explains why these small ectotherm amniotes (the body mass of the largest true salamanders is ≈ 50g) have lifespans similar to that of large endotherm amniotes (e.g., ungulates, large birds) and undergo a marginal actuarial senescence contrary to most vertebrates including humans (Jones et al. 2014).

## Supporting information

Sup_Mat_1

## References

Ackermann, M., Stearns, S. C., Jenal, U. (2003). Senescence in a bacterium with asymmetric division. Science, 300, 1920.

Angelini, C., (2006). Ecologia di popolazione di *Salamandrina perspicillata* (Savi, 1821) (Amphibia, Salamandridae). PhD thesis, Università “la Sapienza” di Roma.

Angelini, C., Antonelli, D., & Utzeri, C. (2010). Capture-mark-recapture analysis reveals survival correlates in *Salamandrina perspicillata* (Savi, 1821). Amphibia-Reptilia, 31, 21–26.

Baudisch, A., & Vaupel, J. W. (2012). Getting to the root of aging. Science, 338, 618–619.

Baudisch, A., Salguero-Gómez, R., Jones, O. R., Wrycza, T., Mbeau-Ache, C., Franco, M., & Colchero, F. (2013). The pace and shape of senescence in angiosperms. Journal of Ecology, 101, 596–606.

Bendik, N. F. (2017). Demographics, reproduction, growth, and abundance of Jollyville Plateau salamanders (*Eurycea tonkawae*). Ecology and Evolution, 7, 5002–5015.

Bielby, J., Mace, G. M., Bininda-Emonds, O. R., Cardillo, M., Gittleman, J. L., Jones, K. E., Orme, C. D. L., & Purvis, A. (2007). The fast-slow continuum in mammalian life history: an empirical reevaluation. The American Naturalist, 169, 748–757.

Cayuela, H., Arsovski, D., Thirion, J. M., Bonnaire, E., Pichenot, J., Boitaud, S., Brison, A.-L., Miaud, C., Joly, P., & Besnard, A. (2016). Contrasting patterns of environmental fluctuation contribute to divergent life histories among amphibian populations. Ecology, 97, 980–991.

Cayuela, H., Joly, P., Schmidt, B. R., Pichenot, J., Bonnaire, E., Priol, P., Peyronel, O., Laville, M., & Besnard, A. (2017). Life history tactics shape amphibians’ demographic responses to the North Atlantic Oscillation. Global Change Biology, 23, 4620–4638.

Cayuela, H., Schmidt, B. R., Weinbach, A., Besnard, A., & Joly, P. (2019). Multiple density-dependent processes shape the dynamics of a spatially structured amphibian population. Journal of Animal Ecology, 88, 164–177.

Choquet, R., Rouan, L., & Pradel, R. (2009). Program E-SURGE: a software application for fitting multievent models. In: Thomson, D. L., Cooch, E. G., Conroy, M. J. (eds) Modeling demographic processes in marked populations. Pp. 845–865, Springer, Boston, US.

Colchero, F., & Clark, J. S. (2012a). Bayesian inference on age-specific survival for censored and truncated data. Journal of Animal Ecology, 81, 139–149.

Colchero, F., Jones, O. R., & Rebke, M. (2012b). BaSTA: an R package for Bayesian estimation of age-specific survival from incomplete mark–recapture/recovery data with covariates. Methods in Ecology and Evolution, 3, 466–470.

Colchero, F., & Schaible, R. (2014). Mortality as a bivariate function of age and size in indeterminate growers. Ecosphere, 5(12), 1–14.

Colchero, F., Jones, O. R., Conde, D. A., Hodgson, D., Zajitschek, F., Schmidt, B. R., Malo, A. F., Alberts, S. C., Becker, P. H., Bouwhuis, S. Bronikowski, A. M., De Vleeschouwer, K. M., Delahay, R. J., Dummermuth, S., Fernández-Duque, E., Frisenvænge, J., Hesselsøe, M., Larson, S., J.-F. Lemaître, McDonald, J., Miller, D. A. W., O’Donnell, C., Packer, C., Raboy, B. E., Reading, C. J., Wapstra, E., Weimerskirch, H., While, G. M., Baudisch, A., Flatt, T., Coulson, T., & Gaillard, J.-M. (2019). The diversity of population responses to environmental change. Ecology Letters, 22, 342–353.

Dammhahn, M., Dingemanse, N. J., Niemelä, P. T., & Réale, D. (2018). Pace-of-life syndromes: a framework for the adaptive integration of behaviour, physiology and life history. 72, 62.

Deelen, J., Beekman, M., Capri, M., Franceschi, C., & Slagboom, P. E. (2013). Identifying the genomic determinants of aging and longevity in human population studies: progress and challenges. Bioessays, 35, 386–396.

Lemaître, J. F., & Gaillard, J. M. (2017). Reproductive senescence: new perspectives in the wild. Biological Reviews, 92, 2182–2199.

Greven, H. & Thiesmeier, B. (1994). Biology of Salamandra and Mertensiella. DGHT, Bonn.

Hamilton, W.D. (1966) Moulding of senescence by natural selection. Journal of Theoretical Biology, 12, 12–45.

Harvey, P. H., & Zammuto, R. M. (1985). Patterns of mortality and age at first reproduction in natural populations of mammals. Nature, 315, 319.

Ivanov, D. K., Escott-Price, V., Ziehm, M., Magwire, M. M., Mackay, T. F., Partridge, L., & Thornton, J. M. (2015). Longevity GWAS using the Drosophila genetic reference panel. Journals of Gerontology Series A: Biomedical Sciences and Medical Sciences, 70, 1470–1478.

Jehle, R., Hoedl, W., & Thonke, A. (1995). Structure and dynamics of central European amphibian populations: a comparison between *Triturus dobrogicus* (Amphibia, Urodela) and *Pelobates fuscus* (Amphibia, Anura). Australian Journal of Ecology, 20, 362–366.

Jones O. R., Gaillard J.-M., Tuljapurkar S., Alho J. S., Armitage K. B., Becker P. H., Bize P., Brommer, J., Charmantier, A., Charpentier, M., Clutton-Brock, T., Dobson, F. S., Festa-Bianchet, M., Gustafsson, L., Jensen, H, Jones, C. G., Lillandt, B. G., Mc Cleery, R., Merilä, J., Neuhaus, P., Nicoll, M. A. C., Norris, K., Oli, M. K., Pemberton, J., Pietiäinen, H., Ringsby, T. H., Roulin, A., Saether, B. E., Setchell, J. M., Sheldon, B. C., Thompson, P. M., Weimerskirch, H., Wickings, E. J., & Coulson, T. (2008). Senescence rates are determined by ranking on the fast–slow life-history continuum. Ecology Letters, 11, 664–673.

Jones, O. R., Scheuerlein, A., Salguero-Gómez, R., Camarda, C. G., Schaible, R., Casper, B. B., Dahlgren, J. P., Ehrlén, J., García, M. B., Menges, E. S., Quintana-Ascencio, P. F., Caswell, H., Baudisch, A., & Vaupel, J. W. (2014). Diversity of ageing across the tree of life. Nature, 505, 169–173.

Jones, O. R., & Vaupel, J. W. (2017). Senescence is not inevitable. Biogerontology, 18, 965–971.

Kirkwood, T. B., & Rose, M. R. (1991). Evolution of senescence: late survival sacrificed for reproduction. Philosophical Transactions of the Royal Society of London. Series B: Biological Sciences, 332, 15–24.

Kirkwood, T. B., & Austad, S. N. (2000). Why do we age?. Nature, 408, 233.

Kirkwood, T. B. (2017). The disposable soma theory. Shefferson, R. P., Jones, O. R., Salguero-Gomez, R. Evolution of senescence in the tree of life. Pp 23–39. Cambridge University Press, Cambridge, UK.

Kopp, M., & Baur, B. (2000). Intra-and inter-litter variation in life-history traits in a population of fire salamanders (*Salamandra salamandra terrestris*). Journal of Zoology, 250, 231–236.

Lemaître, J. F., Berger, V., Bonenfant, C., Douhard, M., Gamelon, M., Plard, F., & Gaillard, J. M. (2015). Early-late life trade-offs and the evolution of ageing in the wild. Proceedings of the Royal Society B: Biological Sciences, 282, 20150209.

Luiselli, L., Andreone, F., Capizzi, D., & Anibaldi, C. (2001). Body size, population structure and fecundity traits of a *Salamandra atra atra* (Amphibia, Urodela, Salamandridae) population from the northeastern Italian Alps. Italian Journal of Zoology, 68, 125–130.

Ma, S., & Gladyshev, V. N. (2017). Molecular signatures of longevity: insights from cross-species comparative studies. Seminars in Cell & Developmental Biology, 70, 90–203.

Martinez, D. E. (1998). Mortality patterns suggest lack of senescence in Hydra. Experimental Gerontology, 33, 217–225.

Miller, D. A., Janzen, F. J., Fellers, G. M., Kleeman, P. M., & Bronikowski, A. M. (2014). Biodemography of ectothermic tetrapods provides insights into the evolution and plasticity of mortality patterns. Sociality, Hierarchy, Health: Comparative Biodemography, National Academies Press, Washington.

Monaghan, P., Charmantier, A., Nussey, D. H., & Ricklefs, R. E. (2008). The evolutionary ecology of senescence. Functional Ecology, 22, 371–378.

Muths, E., Chambert, T., Schmidt, B. R., Miller, D. A. W., Hossack, B. R., Joly, P., Grolet, O., Green, D. M., Pilliod, D. S., Cheylan, M., Fisher, R. N., McCaffery, R. M., Adams, M. J., Palen, W. J., Arntzen, J. W., Garwood, J., Fellers, G., Thirion, J.-M., Besnard, A., & Fisher, R. N. (2017). Heterogeneous responses of temperate-zone amphibian populations to climate change complicates conservation planning. Scientific Reports, 7, 17102.

Nussey, D. H., Froy, H., Lemaitre, J. F., Gaillard, J. M., & Austad, S. N. (2013). Senescence in natural populations of animals: widespread evidence and its implications for bio-gerontology. Ageing Research Reviews, 12, 214–225.

Olgun, K., Miaud, C., & Gautier, P. (2001). Age, growth, and survivorship in the viviparous salamander *Mertensiella luschani* from southwestern Turkey. Canadian Journal of Zoology, 79, 1559–1567.

Oliveira, B. F., São-Pedro, V. A., Santos-Barrera, G., Penone, C., & Costa, G. C. (2017). AmphiBIO, a global database for amphibian ecological traits. Scientific data, 4, 1

Pianka, E. R. (1970). On r-and K-selection. The American Naturalist, 104, 592–597.

Pletcher SD. 1999 Model fitting and hypothesis testing for age-specific mortality data. Journal of Evolutionary Biology, 12, 430–439.

Poss, K. D. (2010). Advances in understanding tissue regenerative capacity and mechanisms in animals. Nature Reviews Genetics, 11, 710.

Pradel, R. (1996). Utilization of capture-mark-recapture for the study of recruitment and population growth rate. Biometrics, 52, 703–709.

Pradel, R. (2005). Multievent: an extension of multistate capture–recapture models to uncertain states. Biometrics, 61, 442–447.

Promislow, D. E., & Harvey, P. H. (1990). Living fast and dying young: A comparative analysis of life-history variation among mammals. Journal of Zoology, 220, 417–437.

Reinhard, S., Renner, S., & Kupfer, A. (2015). Age and fecundity in *Salamandra algira* (Caudata: Salamandridae). Salamandra, 51, 19–24.

Remolina SC, Chang PL, Leips J, Nuzhdin SV, Hughes KA (2012). Genomic basis of aging and life-history evolution in *Drosophila melanogaster*. Evolution, 66, 3390–3403.

Rose, M. R. (1994). Evolutionary biology of aging. Oxford University Press, Oxford, UK.

Rovelli, V., Randi, E., Davoli, F., Macale, D., Bologna, M. A., & Vignoli, L. (2015). She gets many and she chooses the best: polygynandry in *Salamandrina perspicillata* (Amphibia: Salamandridae). Biological Journal of the Linnean Society, 116, 671–683.

Sæther, B. E. (1988). Pattern of covariation between life-history traits of European birds. Nature, 331, 616.

Schmidt, B. R., Feldmann, R., & Schaub, M. (2005). Demographic processes underlying population growth and decline in *Salamandra salamandra*. Conservation Biology, 19, 1149–1156.

Schmidt, B. R., Schaub, M., & Steinfartz, S. (2007). Apparent survival of the salamander *Salamandra salamandra* is low because of high migratory activity. Frontiers in Zoology, 4, 19.

Schmidt, B. R., Hödl, W., & Schaub, M. (2012). From metamorphosis to maturity in complex life cycles: equal performance of different juvenile life history pathways. Ecology, 93, 657–667.

Seifert, A. W., & Voss, S. R. (2013). Revisiting the relationship between regenerative ability and aging. BMC Biology, 11, 2.

Sinsch, U., Böcking, H., Leskovar, C., Öz, M., & Veith, M. (2017). Demography and lifetime growth patterns in viviparous salamanders (genus Lyciasalamandra): Living underground attenuates interspecific variation. Zoologischer Anzeiger, 269, 48–56.

Siler W. 1979 A competing-risk model for animal mortality. Ecology, 60, 750–757.

Sharon, R., Degani, G., & Warburg, M. R. (1997). Oogenesis and the ovarian cycle in *Salamandra salamandra infraimmaculata* Mertens (Amphibia; Urodela; Salamandridae) in fringe areas of the taxon’s distribution. Journal of morphology, 231, 149–160.

Spiegelhalter, D. J., Best, N. G., Carlin, B. P., & Van Der Linde, A. (2002). Bayesian measures of model complexity and fit. Journal of the Royal Statistical Society: Series B, 64, 583–639.

Stearns, S. C. (1976). Life-history tactics: a review of the ideas. The Quarterly Review of Biology, 51, 3–47.

Vaupel, J. W., Baudisch, A., Dölling, M., Roach, D. A., & Gampe, J. (2004). The case for negative senescence. Theoretical Population Biology, 65, 339–351.

Velo-Antón, G., Zamudio, K. R., & Cordero-Rivera, A. (2012). Genetic drift and rapid evolution of viviparity in insular fire salamanders (*Salamandra salamandra*). Heredity, 108, 410.

Warburg, M. R. (2007). Longevity in *Salamandra infraimmaculata* from Israel with a partial review of life expectancy in urodeles. Salamandra, 43, 21.

Weisrock, D. W., Papenfuss, T. J., Macey, J. R., Litvinchuk, S. N., Polymeni, R., Ugurtas, I. H., Zhao, E., Jowkar, H., & Larson, A. (2006). A molecular assessment of phylogenetic relationships and lineage accumulation rates within the family Salamandridae (Amphibia, Caudata). Molecular Phylogenetics and Evolution, 41, 368–383.

Wells, K. D. (2010). The ecology and behavior of amphibians. University of Chicago Press.

Wright, J., Bolstad, G. H., Araya-Ajoy, Y. G., & Dingemanse, N. J. (2019). Life-history evolution under fluctuating density-dependent selection and the adaptive alignment of pace-of-life syndromes. Biological Reviews, 94(1), 230–247.

Yokoyama, H. (2008). Initiation of limb regeneration: the critical steps for regenerative capacity. Development, Growth & Differentiation, 50, 13–22.

