## Supplementary material for "Slow life-history strategies are associated with negligible actuarial senescence in western Palearctic salamanders": Sup_Mat_1

**Table S1**. Information about age-dependent capture-recapture data in four populations of *Lyciasalamandra fazilae*, *Salamandrina perspecillata*, and *Salamandra salamandra* (two populations).

| Parameter | *L. fazilae* | *S. perspecillata* | *S. salamandra 1* | *S. salamandra 2* |
| --- | --- | --- | --- | --- |
| Number of individuals | 133 | 911 | 304 | 376 |
| Number with known birth year | 67 | 55 | 0 | 0 |
| Number with known death year | 5 | 0 | 0 | 0 |
| Total number of captures | 179 | 1629 | 609 | 1474 |
| Earliest detection time | 1999 | 1998 | 2008 | 1965 |
| Latest detection time | 2009 | 2006 | 2015 | 1985 |
| Earliest recorded birth year | 1990 | 1994 | 0 | 0 |
| Latest recorded birth year | 1997 | 2002 | 0 | 0 |
| Earliest recorded death year | 2000 | 0 | 0 | 0 |
| Latest recorded death year | 2003 | 0 | 0 | 0 |
